# Oncolytic virus treatment differentially affects the CD56^dim^ and CD56^bright^ NK cell subsets *in vivo* and regulates a spectrum of human NK cell activity

**DOI:** 10.1101/2020.12.16.423062

**Authors:** Michelle Wantoch, Erica B. Wilson, Alastair P. Droop, Sarah L. Phillips, Matt Coffey, Yasser M. El-Sherbiny, Tim D. Holmes, Alan A. Melcher, Laura F. Wetherill, Graham P. Cook

## Abstract

Natural killer (NK) cells protect against intracellular infection and cancer. These properties are exploited in oncolytic virus (OV) therapy, where anti-viral responses enhance anti-tumour immunity. We have analysed the mechanism by which reovirus, an oncolytic dsRNA virus, modulates human NK cell activity. Reovirus activates NK cells in a type I interferon (IFN-I) dependent manner, resulting in STAT1 and STAT4 signalling in both CD56^dim^ and CD56^bright^ NK cell subsets. Gene expression profiling revealed the dominance of IFN-I responses and identified induction of genes associated with NK cell cytotoxicity and cell cycle progression, with distinct responses in the CD56^dim^ and CD56^bright^ subsets. However, reovirus treatment, acting via IFN-I, inhibited NK cell proliferative responses to IL-15 and was associated with reduced AKT signalling. *In vivo*, human CD56^dim^ and CD56^bright^ NK cells responded with similar kinetics to reovirus treatment, but CD56^bright^ NK cells were transiently lost from the peripheral circulation at the peak of the IFN-I response, suggestive of their redistribution to secondary lymphoid tissue. These results show that reovirus modulates a spectrum of NK cell activity *in vivo*, encompassing direct action on tumour cells and the regulation of adaptive immunity. Such activity is likely to mirror NK cell responses to natural viral infection.

## 1. Introduction

Natural killer (NK) cells are innate lymphoid cells (ILCs) with two broad functions, they detect and destroy infected cells and tumour cells and they provide signals for the initiation of adaptive immunity (1, 2). Humans with NK cell defects are highly susceptible to viral infection (3, 4) and many viruses, herpesviruses and poxviruses in particular, encode numerous gene products that mediate evasion of NK cells (5, 6). Indeed, battles between viruses and NK cells act as a major driver of both viral and NK cell evolution (7, 8). Mouse and human studies have confirmed the importance of NK cells in anti-tumour immunity (9) and the ability of NK cells to respond to both viral infection and cancer makes them a key mediator of the anti-tumour activity of oncolytic viruses.

Oncolytic viruses (OV) are an emerging group of therapeutic agents (10–13). The therapeutic basis of OV activity was initially believed to result from their preferential replication in tumour cells, resulting in direct lysis (14–16). However, it is now widely accepted that lytic activity is accompanied by the stimulation of anti-tumour immunity, with NK cells playing a key role in their action. Depletion of NK cells (or NK cell activity) from tumour-bearing mice reduces the therapeutic effect of several OV (17–20) and human NK cells rapidly respond to oncolytic reovirus treatment *in vivo* (21). Several studies have shown that human NK cell activation by OV is mediated by IFN-I and that this results in enhanced cytotoxicity against tumour cells (22–24). However, the action of OV on human NK cells is not well defined. Studies to date have analysed OV action on bulk NK cell activity, with a focus on NK cell cytotoxicity. However, human NK cells exist in two major subsets; CD56^dim^ NK cells predominate in the blood, they express CD16 and have strong cytotoxic activity, whereas the CD56^bright^ NK cell subset is less cytotoxic, have low (or no) CD16 expression and predominate in the secondary lymphoid tissue (SLT) (25–27). Here we have combined *in vitro* studies with the analysis of a reovirus clinical trial to gain insight into the action of OV on human NK cells. Our results demonstrate that oncolytic reovirus differentially affects the CD56^dim^ and CD56^bright^ NK cell subsets, modulating direct anti-tumour effects as well as responses that likely regulate adaptive immunity.

## 2. Materials and methods

### 2.1 Cells

Peripheral Blood Mononuclear Cells (PBMC) were obtained from waste apheresis cones from healthy donors via NHS Blood and Transplant (UK). On the day of donation, PBMCs were separated by density gradient centrifugation using Lymphoprep (Axis-Shield) and cultured in Roswell Park Memorial Institute medium (RPMI) 1640 (Sigma) +10% foetal calf serum (FCS) at a concentration of 2×10^6^ cells/ml at 37°C, 5% CO_2_. Natural killer (NK) cells were separated from PBMC, either on the day of donation or after the time specified, by negative immunomagnetic selection (Miltenyi Biotec). NK cells were cultured in Dulbecco’s Modified Eagle Medium (DMEM) supplemented with 10% human AB serum (Gemini Bio-Products) and 10% FCS at 37°C, 5% CO_2_ unless otherwise specified.

### 2.2 Reovirus and the clinical trial

Clinical grade reovirus (Pelareorep; formerly known as Reolysin) was provided by Oncolytics Inc. (Canada) and viral titres determined by routine plaque assays on L929 cells. Overnight cultures of PBMC were treated with reovirus at a multiplicity of infection (MOI) of 1 (unless otherwise stated). For the trial, ten patients with colorectal liver metastases were treated with intravenous reovirus prior to surgical resection of their tumour. Blood samples from reovirus-treated patients were analysed immediately after collection. The clinical study was undertaken at the Leeds Cancer Centre following full ethical and regulatory approval. Patients were enrolled into the trial (and provided blood samples) following informed consent. The patient group and the clinical trial, including dose and scheduling of the reovirus treatment has been described previously (21, 28).

### 2.3 Cytokine treatment

PBMC or NK cells were cultured overnight before treating with purified IFN-α (Sigma) or recombinant IFN-α (Miltenyi Biotec), recombinant IL-12 (R&D systems and Peprotech) or recombinant IL-15 (Miltenyi Biotec) as described in the text and figure legends. For IFN-I neutralisation, a cocktail of anti-human interferon α/β receptor chain 2 antibody (clone MMHAR-2), anti-human interferon-α (sheep polyclonal) and anti-human interferon-β (sheep polyclonal; all from PBL Assay Science) or a control cocktail of mouse IgG2a (BioLegend) and sheep serum (Sigma) was used, as described previously (21).

### 2.4 Immunoblotting and ELISA assays

For immunoblotting, cells were washed with PBS and lysed in RIPA buffer with added protease and phosphatase inhibitors (Roche). Samples were sonicated, diluted in Laemmli sample buffer and separated by SDS-PAGE, before transfer to PVDF membrane. Membranes were probed with antibodies listed in Supplementary Table 1 and a secondary antibody conjugated to horseradish peroxidase, before developing using enhanced chemiluminesence substrate. For IFN-I ELISA, 96 well plates were coated with a mixture of antibodies against IFN-α (Mabtech, MT1/3/5) overnight at 4°C, and blocked with PBS supplemented with 10% FCS. Samples along with recombinant IFN-α2 (Miltenyi), for construction of the standard curve, were added in triplicate and incubated overnight at 4°C. The plate was washed and IFN-I assayed using a mixture of biotinylated detection antibodies against IFN-α (Mabtech, MT2/4/6). Avidin-conjugated alkaline phosphatase (Sigma, ExtrAvidin), and p-Nitrophenyl phosphate substrate (Sigma, SigmaFast tablets) were used to develop the ELISA. Absorbance was read at 405nm on a Multiskan EX plate reader (Thermo Fisher).

### 2.5 Flow cytometry

For all flow cytometry experiments, staining buffer (PBS+2% FCS+0.09% sodium azide) was used for washing and staining steps. Isotype matched control antibodies were used in all experiments and a gate set whereby 2% of isotype control antibody stained cells were positive; for test antibodies, cells staining within this gate were assessed as positive. Cells were analysed on a LSRII flow cytometer (BD Biosciences) or a Cytoflex cytometer (Beckman Coulter). For cell sorting, an Influx cell sorter (BD Biosciences) was used. All antibodies used in flow cytometry experiments are listed in Supplementary Table 1. NK cells within PBMC were identified as the CD56^+^CD3^neg^ population (Supplementary Figure S1).

### 2.6 Intracellular staining

For granzyme B, cells were first stained for surface proteins, fixed in Cytofix buffer (BD Biosciences), washed and incubated in saponin buffer (staining buffer + 0.1% saponin), and stained in saponin buffer plus anti-granzyme B antibody (or a matched isotype control). Cells were resuspended in 0.5% paraformaldehyde (in staining buffer) prior to analysis. For other intracellular proteins, cells were stained for surface proteins if required and then fixed in Cytofix fixation buffer (BD Biosciences), according to the manufacturer’s instructions. For time-course experiments, fixed cells were stored at 4°C and all cells within an experiment were stained at the same time. Samples were then resuspended in Permeabilisation Buffer III (BD Biosciences) and permeabilised on ice for 30 minutes, followed by staining, washing and analysis.

### 2.7 NK cell degranulation assay

PBMC were co-cultured with K562 cells in a 96 well, round bottomed cell culture plate (Corning), at an effector:target ratio of 10:1. After 1 hour of culture, GolgiStop (BD Biosciences) was added and cells were cultured for a further 5 hours before staining for NK cell markers and cell surface CD107a.

### 2.8 Proliferation and cell cycle profiling

For CFSE labelling, cells were labelled by suspension in warm PBS, containing 2μM CFDA-SE (Invitrogen) and incubation for 10 minutes at 37°C. The labelling reaction was quenched with an equal volume of warm FCS and analysed by flow cytometry. For propidium iodide (PI) staining, washed cells were resuspended in ice cold 70% ethanol at 1×10^6^ cells/ml with vortexing. Samples were fixed on ice for 30 minutes then stored at −20°C. For staining, fixed cells were washed with 2ml stain buffer, centrifuging at 600*xg*. Samples (0.5–1×10^6^ cells) were resuspended in 50μl of 100μg/ml RNase A (Qiagen). PI (Life Technologies) at 16.6μg/ml in staining buffer was added directly to samples in RNase A (final density of 10^6^ cells in 650μl). Samples were incubated at room temperature for 10 minutes and analysed using an LSRII flow cytometer (BD Biosciences), on the lowest speed setting. Cell cycle profiling was performed using Modfit (Verity software) according to the manufacturer’s recommendations.

### 2.9 Gene expression profiling and data analysis

PBMC isolated from five healthy donors were cultured with or without 1 MOI reovirus for 48 hours. NK cells were isolated by immunomagnetic selection and resuspended in RNAprotect Cell Reagent (Qiagen); RNA was extracted with the RNeasy mini kit (Qiagen), according to the manufacturer’s instructions. Contaminating DNA was removed by on-column DNase digestion and RNA integrity was checked using an Agilent 2100 Bioanalyzer (Agilent Technologies). Total RNA was amplified, sense strand cDNA synthesised and labelled using the GeneChip™ WT PLUS Reagent Kit (Applied Biosystems, Thermo Fisher Scientific). Labelled cDNA was hybridised to an Affymetrix GeneChip® Human Transcriptome Array 2.0 (Applied Biosystems, Thermo Fisher Scientific). Raw intensity files (CEL files) for all conditions were processed with the Expression Console software (Affymetrix) using the Signal Space Transformation-Robust Multi-Chip Analysis (SST-RMA) algorithm. Normalised signal values were analysed with the Transcriptome Analysis Console (TAC, Affymetrix) software, to identify statistically significant differences between conditions. Following manufacturer guidelines, TAC software was used to run paired ANOVA tests and false discovery rate (FDR) prediction. Differentially expressed genes were defined as >1.5 fold up or downregulated with FDR<0.05. Gene set enrichment analysis was performed using the tools provided by Enrichr (29, 30); available at https://amp.pharm.mssm.edu/Enrichr/. Interferon regulation of genes was analysed using the interferome database (31); available at http://www.interferome.org/interferome/home.jspx. We used v2.01 of the database and restricted our analysis to human genes regulated (>1.5 fold change) by IFN-I in haematopoietic cells, using the filters provided. To analyse intersection of datasets we used the Venn diagram drawing tool available at; http://bioinformatics.psb.ugent.be/webtools/Venn/. The microarray data is available at the EMBL-EBI Array Express repository (https://www.ebi.ac.uk/arrayexpress/) with the accession number E-MTAB-9826.

### 2.10 Quantitative RT-PCR

Cells were resuspended and stored in RNAprotect cell reagent (Qiagen). RNA was extracted with the RNeasy mini kit (Qiagen) and contaminating DNA removed by on column DNase digestion, with the RNase-Free DNase Set (Qiagen). cDNA was synthesised using random primers (New England Biolabs) and the Superscript III reverse transcriptase kit (Invitrogen). qPCR amplification was performed with either Taqman or SYBR Green reagents. For both, 10ng cDNA was used together with assay specific primers (Supplementary Table 2), and either PowerUp™ SYBR™ Green Master Mix (for SYBR Green method) or Taqman gene expression Mastermix (both Applied Biosystems). Amplification was carried out in a 7500 Real Time PCR machine or a QuantStudio 5 machine (Applied Biosystems). Reactions were performed in triplicate and all replicate values were within 0.5 cycles. Fold change gene expression was calculated by the ΔΔCt method (32), normalizing to ABL1 as the housekeeping gene.

### 2.11 Statistical testing

Tests (identified in the figure legends) were performed using Prism (GraphPad Software).

## 3. Results

Several cytokines are implicated in virus-dependent activation of NK cell activity. In particular, IFN-I, IL-12 and IL-15 have established roles in viral infection and NK activation (33, 34). Signal transduction pathways for these cytokines overlap, but they utilise characteristic JAK/STAT pathways that can be used to indicate their activity. Current models of cytokine signalling indicate that IFN-I signals predominantly via STAT1 and STAT4, IL-12 via STAT4 and IL-15 via STAT5 (35–38); these patterns were verified using immunoblotting of lysates from purified human NK cells treated with recombinant cytokines (Figure 1A). Furthermore, this pattern of STAT phosphorylation was also observed using intracellular staining and flow cytometry of NK cells present in cytokine-treated PBMC (Figure 1B and Supplementary Figure S2). Combining intracellular staining of phosphorylated STAT molecules with detection of cell surface markers allowed the responses of CD56^bright^ and CD56^dim^ NK cells to be analysed separately. The two subsets responded similarly to cytokines, the major difference being that CD56^bright^ NK cells responded to IL-12 more strongly than their CD56^dim^ counterparts (Figure 1B), as shown previously (39). In a timecourse experiment, IFN-I treatment resulted in rapid and transient phosphorylation of STAT1 and STAT4, whereas IL-12 and IL-15 led to sustained STAT4 and STAT5 responses respectively, again with similar responses in both CD56^dim^ and CD56^bright^ NK cells (Figure 1C).

**Figure 1:**
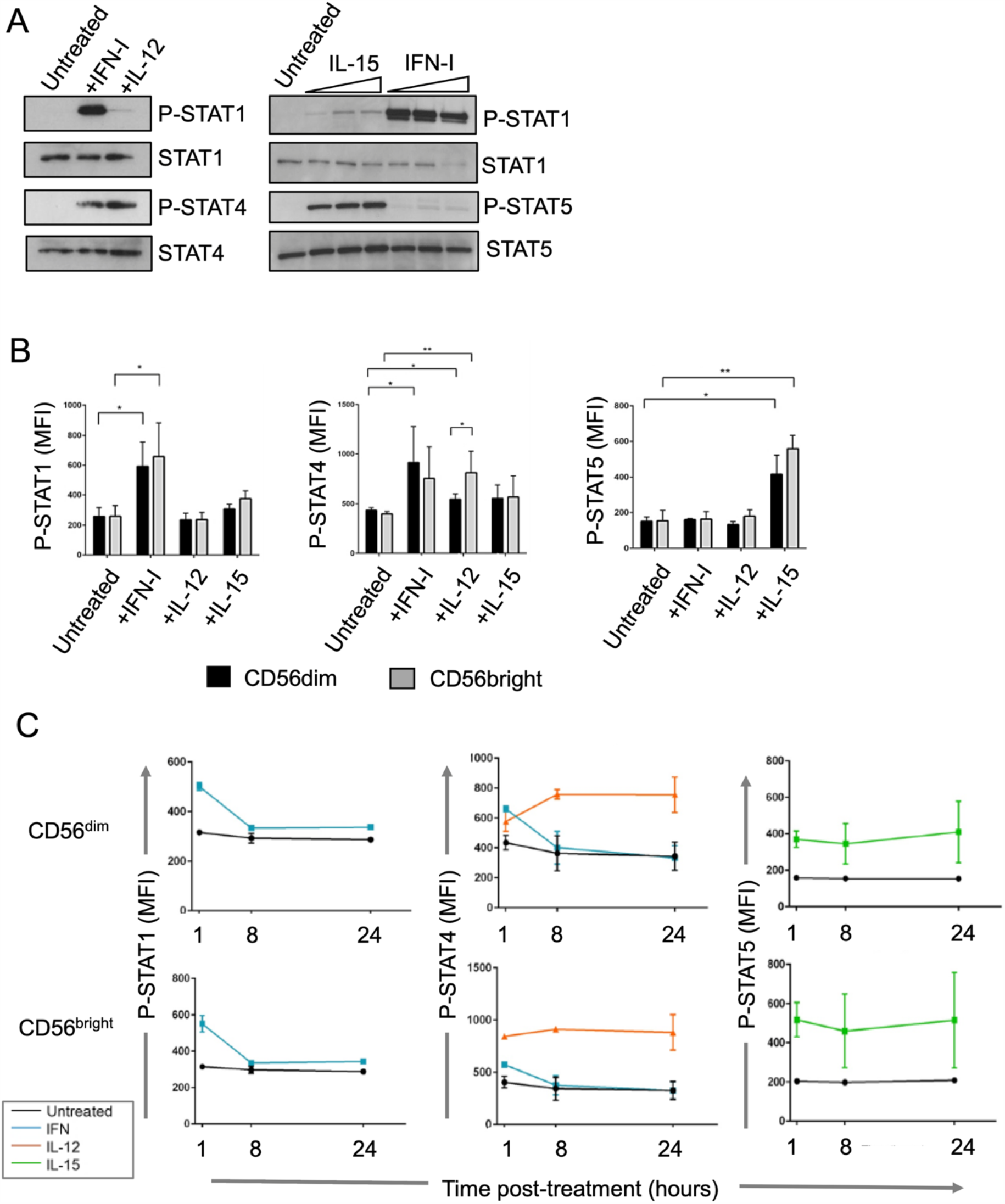
STAT phosphorylation dynamics in cytokine treated CD56^dim^ and CD56^bright^ NK cells. **A)** STAT phosphorylation in purified, total human NK cells. Cells were treated with 100 IU/mL IFN-I, 10 ng/mL IL-12 or increasing concentrations of IL-15 and IFN-I for 1 hour, and blotted for phosphorylated or total STAT1, STAT4 and STAT5 as indicated. **B)** STAT phosphorylation in CD56^bright^ and CD56^dim^ NK cells (detected by intracellular staining and flow cytometry) in PBMC treated with 100 IU/ml IFN-α, 10 ng/ml IL-12 or 50 IU/ml IL-15 for 1 hour. The flow cytometry gating strategy is shown in Supplementary Figure 1A. Graphs show median fluorescence intensities (MFI), with mean and standard deviation from three donors, from CD56^dim^ (black bars) and CD56^bright^ NK cells (grey bars) as indicated. Data was analysed by one-way repeated measures ANOVA, followed by Dunnet’s multiple comparison test; *p<0.05, **p<0.01. **C)** Time course of STAT phosphorylation in CD56^bright^ and CD56^dim^ NK cells in PBMC treated with cytokines as in panel B, fixed at 1, 8 and 24 hours and analysed by intracellular staining and flow cytometry. Graphs show mean MFI and standard deviation from two donors. The colour key indicates cytokine treatment.

We then analysed the responses of NK cells within reovirus-treated PBMC, modelling the intravenous delivery of reovirus used in our clinical trials of this agent in colorectal cancer and glioblastoma (28, 40). We treated PBMC at a multiplicity of infection (MOI) of 1 (approximating the dose used in the clinical trials) and analysed STAT phosphorylation in NK cells at 8, 24 and 48 hours post-treatment, reasoning that cytokines induced during treatment would take time to accumulate. For both the CD56^dim^ and CD56^bright^ NK cell subsets, the level of phosphorylation of STAT1 and STAT4 seen with reovirus treatment of PBMC (Figure 2A) was similar to those seen with IFN-I and IL-12 treatment respectively (Figure 1). The level of STAT5 phosphorylation observed in the presence of reovirus was statistically significant in the CD56^bright^ NK cells (Figure 2A), but was a very small effect compared to that observed with both the CD56^bright^ and CD56^dim^ NK cell subsets treated with IL-15 (Figure 1A). These data are consistent with IFN-I induced signalling in both CD56^bright^ and CD56^dim^ NK cells. The duration of STAT1 and STAT4 phosphorylation differed from the responses obtained using purified cytokines, with STAT1 phosphorylation being maintained (and STAT4 phosphorylation declining) in response to reovirus (Figure 2A). As expected, IFN-I was readily detectable in the supernatants of reovirus treated PBMC (Figure 2B). Reovirus conditioned media (rCM) from PBMC treated with reovirus for 24 hours was collected, filtered to remove virus particles and added to purified NK cells in the presence or absence of anti-IFN antibodies; rCM induced cell surface expression of CD69, a marker of NK cell activation, and tetherin, an IFN induced anti-viral protein, on purified NK cells in an IFN-I dependent manner (Figure 2C). These data suggest that the IFN-I dependent NK cell activation observed in response to reovirus is associated with the direct action of IFN-I on the NK cells themselves.

**Figure 2:**
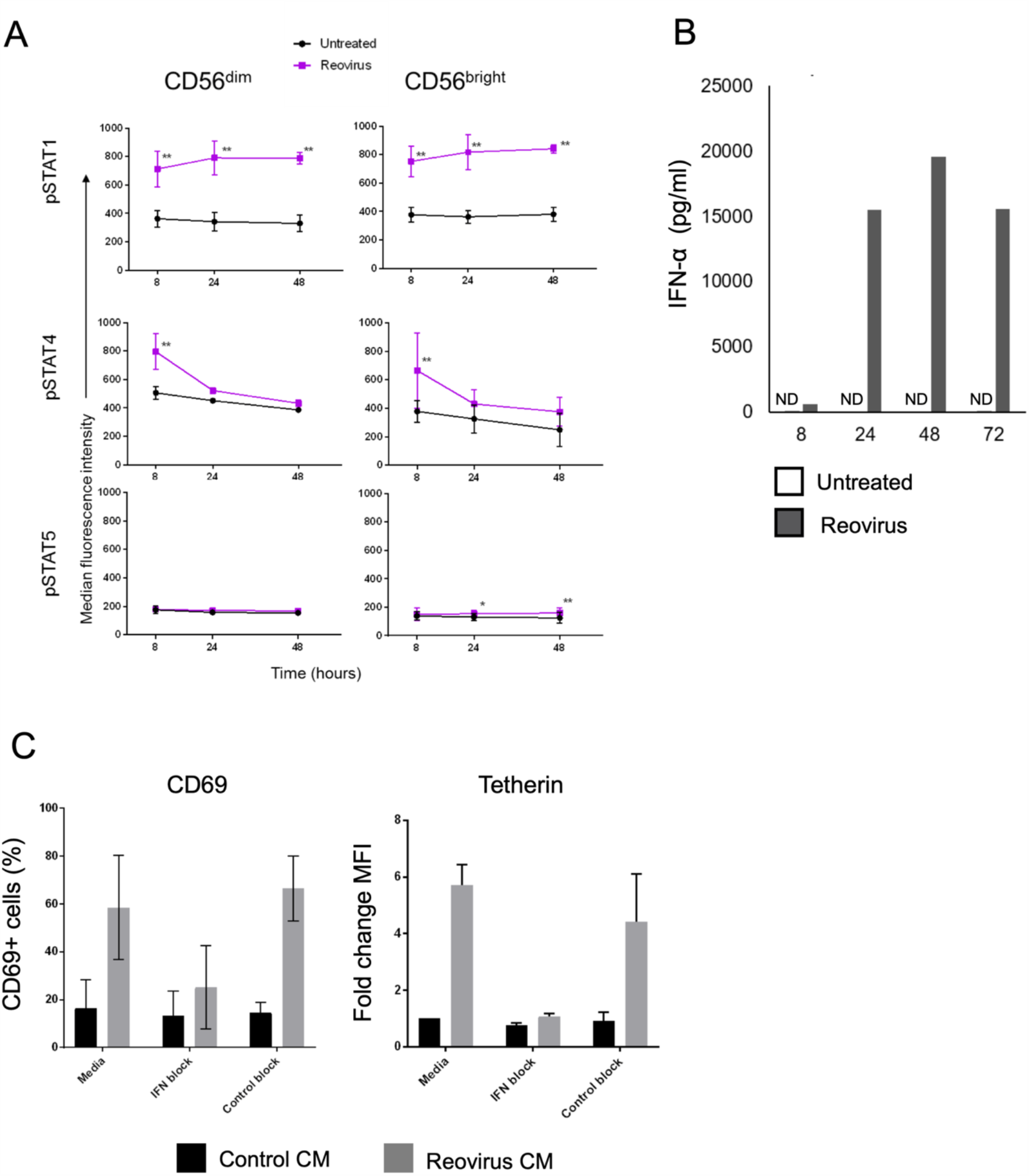
STAT phosphorylation and NK cell activation following reovirus treatment. **A)** STAT phosphorylation in CD56^bright^ and CD56^dim^ NK cells (detected by intracellular staining and flow cytometry) in PBMC cultured without virus (untreated; black line) or with 1 MOI reovirus (purple line), for 8, 24 and 48 hours. Graphs show mean MFI and standard deviation from three donors. Data was analysed by two-way repeated measures ANOVA, followed by Sidak multiple comparisons test. *p<0.05 **p<0.01. **B)**IFN-α production by PBMC following treatment with 1 MOI reovirus (or left untreated). Supernatants was collected and analysed by ELISA. IFN levels were not detectable (ND) in the untreated cells. **C)** NK cell activation (as indicated by cell surface expression of CD69 and tetherin). PBMC (from one donor) were left untreated or treated with reovirus for 24 hours and the conditioned media (CM) filtered to remove viruses. CM was added to purified NK cells in the presence of an IFN-I blocking antibody cocktail (IFN block), a control blocking cocktail (control block) or no added antibody (media). CM from untreated PBMC was used a control. After 48 hours, the NK cell surface expression of CD69 and tetherin was measured by flow cytometry. Data is from control CM or CM from reovirus-treated PBMC from a single donor, applied to three NK cell donors. The y-axis shows the percentage of CD69 expressing cells (left panel) or the fold change in MFI of tetherin relative to control CM and no added antibody treatment (right panel), due to constitutive low level expression of this marker on unstimulated NK cells (20).

To further explore the responses of NK cells to reovirus treatment, we performed gene expression profiling. NK cells were isolated from untreated and reovirus treated PBMC and differentially regulated genes were identified. The NK cells from five healthy donors upregulated cell surface expression of CD69, consistent with NK cell activation following virus treatment (Supplementary Figure S3). Differentially regulated transcripts were detected by 2777 probes, representing some 1742 genes, with an approximately equal number of genes induced and repressed by reovirus treatment (Figure 3A and Supplementary Table 3). Amongst the upregulated genes were IFN stimulated genes (ISGs), including IFI44L and IFIT1, which were also induced in NK cells following reovirus treatment of cancer patients (21). Similarly, BST2 (encoding tetherin) and CD69 were induced at the mRNA level, consistent with detection of their protein products at the cell surface following reovirus treatment *in vivo* (21). Genes whose expression was downregulated by reovirus treatment included FCGR3A (encoding CD16), the NK cell receptors KLRB1 (NKR-P1; CD161) and NCR3 (NKp30; CD337) and the sphingosine-1-phosphate receptor, S1PR1. We used gene set enrichment analysis (GSEA) to analyse the effects of reovirus on NK cells in more detail, exploiting the multiple tools available via the Enrichr suite (29, 30). As expected, GSEA identified IFN regulated pathways as the most highly enriched amongst the 1742 differentially expressed genes, but also highlighted enrichment of genes regulating the cell cycle (Figure 3B and Supplementary Table 4). Accordingly, transcription factors associated with the differentially expressed genes included mediators of IFN and inflammatory responses, as well as regulators of the cell cycle (Figure 3C) (41, 42).

**Figure 3:**
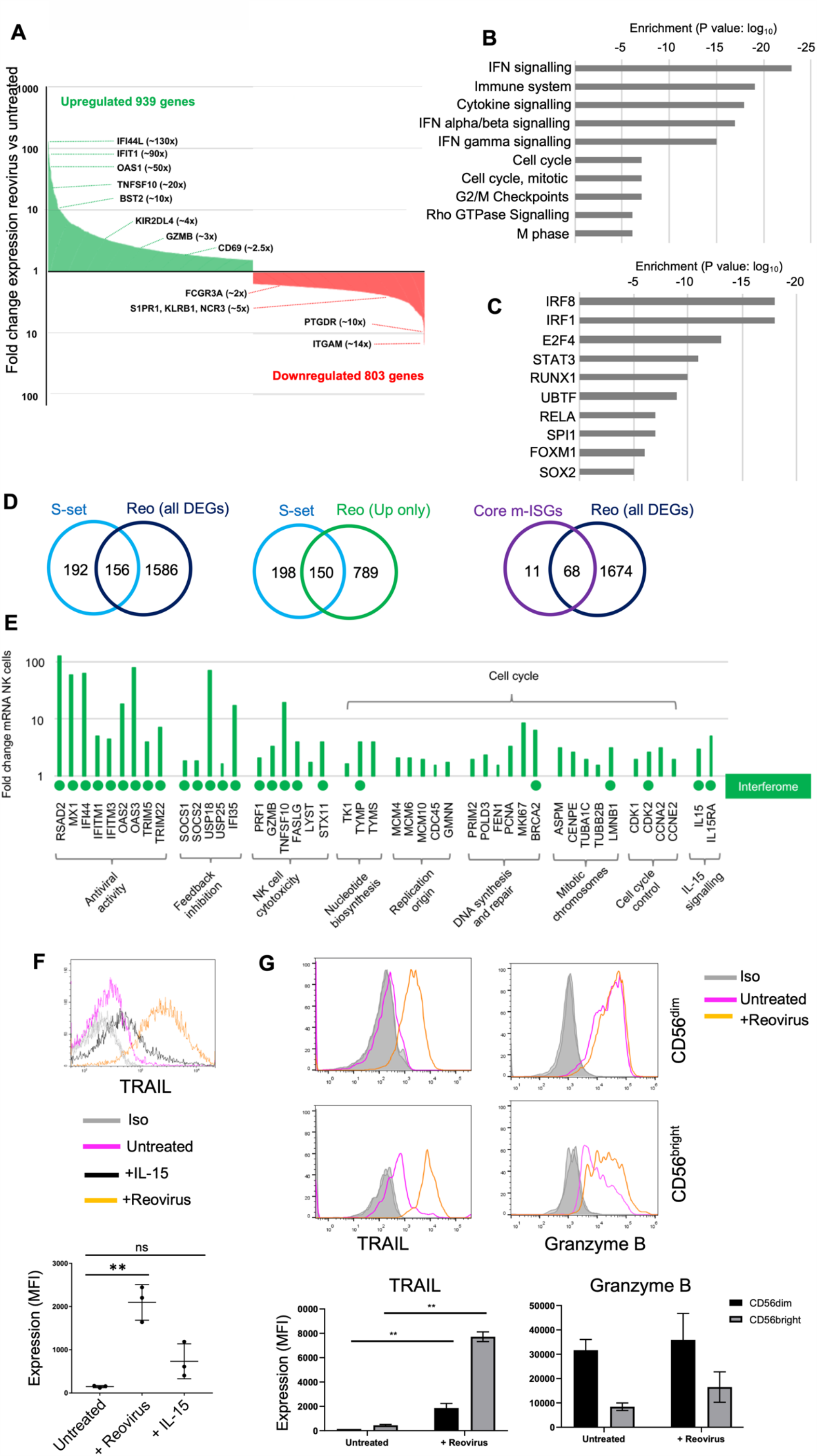
Gene expression profiling of NK cells following reovirus treatment. **A)** Differentially expressed genes in total human NK cells following reovirus treatment. Genes that were up (green columns) or downregulated (red columns) together with examples and fold change are indicated. Data underlying this graph is shown in Supplementary Table 3. **B)** The top ten pathways identified by gene set enrichment analysis of differentially expressed genes. Data were generated using Enrichr and the output from Reactome 2016; the adjusted P value of enrichment is shown. Full data is provided in Supplementary Table 4. **C)** The top ten transcription factors associated with differentially expressed genes. Data were generated using Enrichr and the output from ENCODE and ChEA Consensus Transcription factors from ChIP-X; the adjusted P value of enrichment is shown. Full data are provided in Supplementary Table 5. **D)** Venn diagrams showing the overlap of the differentially expressed genes (DEGs) from NK cells following reovirus treatment (Reo) with the interferon stimulated genes listed in Schoggins et al (41; S-Set) or the core mammalian ISGs identified by Shaw et al (42; Core m-ISGs). Overlaps were determined for all reovirus DEGs (left and right panels) or the upregulated genes only (centre panel). **E)** Expression of genes from selected pathways. The graph shows the fold change in gene expression in NK cells following reovirus treatment, with genes and pathways indicated. A green circle below the x-axis indicates that the gene is induced by IFN-I in haematopoietic cells, as determined using the Interferome database (30). **F)** Induction of TRAIL following reovirus treatment. The top panel shows TRAIL expression from a single representative donor, with NK cells from untreated, IL-15 or reovirus-treated PBMC along with the isotype control stain as indicated. The lower graph shows the MFI of expression from three separate donors. Data were analysed by a repeated measures one-way ANOVA, with Tukey’s multiple comparison test; *p<0.05, **p< 0.01. **G)** TRAIL and granzyme B expression by CD56^dim^ and CD56^bright^ NK cells following reovirus treatment. The top panel shows data from a single representative donor, with the different treatments (and isotype control antibody) indicated along with the NK cell subset analysed via gating. The lower graphs show data from three separate donors, analyse as in panel F; *p<0.05. **p< 0.01.

We analysed the NK cell response to reovirus treatment in more detail. Schoggins et al screened 348 previously identified ISGs for their antiviral effects (43). Of these 348 genes, we found 156 to be differentially regulated in NK cells, with 150 of these 156 genes being upregulated in NK cells during reovirus treatment of PBMC (Figure 3D). In addition, we compared the genes upregulated in NK cells with a set of 79 ISGs induced across nine species of mammals, termed core mammalian ISGs (44); four fifths of these core ISGs, encoding anti-viral and other IFN-I regulated functions were upregulated in the human NK cells following reovirus treatment of PBMC (Figure 3D).

We previously showed that reovirus activates NK cell cytotoxicity in an IFN-I dependent manner (23) and the expression profiling identified PRF1 (encoding perforin), GZMB (granzyme B), TNFSF10 (TRAIL), FASLG (Fas ligand) and two genes, LYST (lysosomal trafficking regulator) and STX11 (syntaxin 11), critical to the biosynthesis and exocytosis of cytotoxic granules (45), as induced by reovirus treatment (Figure 3E). The 20-fold induction of TNFSF10 mRNA was mirrored by strong induction of TRAIL on the NK cell surface, an effect not seen with IL-15 treatment (Figure 3F). We further analysed TRAIL expression on CD56^bright^ and CD56^dim^ NK cells. Without stimulation, TRAIL was expressed at low levels on CD56^bright^ NK cells and was undetectable on the CD56^dim^ NK cells. However, reovirus treatment induced cell surface TRAIL expression on the both subsets (Figure 3G). For the granule-mediated cytotoxic pathway, NK cells showed enhanced degranulation in response to tumour target cells (Supplementary Figure S4), confirming our previous data (21). We analysed intracellular granzyme B levels using flow cytometry and found that the cytotoxic CD56^dim^ NK cell subsets showed little change in granzyme B content, whereas the CD56^bright^ NK cells, which have low cytotoxic activity under resting conditions, did induce expression of granzyme B upon reovirus treatment (Figure 3G). Treatment with IFN-I itself induced a modest increase in granzyme B in purified NK cells and significantly enhanced granule exocytosis, albeit less effectively than IL-15 treatment (Supplementary Figure S4).

The action of IFN-I includes the induction of feedback inhibitory pathways which limit responses (46). In our previous clinical trial, most patients received multiple infusions of reovirus yet only demonstrated NK cell activation after the first virus infusion (21). We speculated that this was due, at least in part, to the induction of feedback inhibitory mechanisms that limit IFN-I responses. We analysed the gene expression profiling for induction of candidate feedback inhibitors and found that the IFN-I signalling antagonists, SOCS1, SOCS2, USP18, USP25 and IFI35 were all induced in NK cells following reovirus treatment of PBMC (Figure 3E).

Our GSEA also revealed the induction of genes associated with cell cycle progression (Figure 3B and E; Supplementary Table 4). This suggested that reovirus treatment of PBMC might stimulate NK cell proliferation. Indeed, studies performed in mice suggest that IFN-I responses induce the expression of IL-15 (e.g. in dendritic cells) and that this, in turn, activates NK cell cytotoxicity and proliferation (33, 34). Gene expression profiling identified induction of the IL-15 receptor α chain gene (IL15RA) in NK cells, as well as increased expression of IL15 itself (Figure 3E). The transcriptional programme downstream of reovirus treatment is clearly dominated by IFN-I responses. However, a variety of stimuli were expected to contribute to the patterns of gene expression observed in NK cells from reovirus treated PBMC. We used the interferome database (31) to assess induction of the 45 genes in Figure 3E in response to IFN-I treatment; IFN-I induces genes encoding antiviral function, feedback inhibition of IFN-I responses, NK cell cytotoxicity and IL-15/IL-15Rα, but the majority of the cell cycle functions were not induced by IFN-I in haematopoietic cells (Figure 3E).

We investigated the induction of cell cycle characteristics in more detail. Reovirus-mediated induction of genes encoding positive regulators of the cell cycle (MCM4, CDK2 and CCNB1) was confirmed using qRT-PCR (Figure 4A and Supplementary Figure S5). Induction of MCM4 mRNA was associated with a significant increase in MCM4 protein 48 hrs post-reovirus treatment (Figure 4B and C). Cell sorting showed that both CD56^bright^ and CD56^dim^ NK cells from reovirus treated PBMC induced the expression of IFIT1 mRNA, whereas MCM4 and IFNG were preferentially induced in the CD56^bright^ population (Figure 4D). The induction of cell cycle-related molecules is suggestive of NK cell proliferation. Markers of proliferating cells include Proliferating Cell Nuclear Antigen (PCNA) and Ki67 and the genes encoding these molecules were induced in NK cells (Figure 3E). At the protein level, the reovirus-mediated induction of PCNA in NK cells was much weaker than that observed with IL-15. Nevertheless, reovirus did induce PCNA expression in both CD56^dim^ and CD56^bright^ NK cells (albeit not reaching statistical significance), with greater induction in the latter subset (Figure 4E and F). Similar experiments analysing Ki67 expression failed to show induction of this marker in either NK cell subset following reovirus treatment, although expression in response to IL-15 was observed (Supplementary Figure S6). However, despite the proliferative gene expression signature, reovirus treatment of PBMC did not induce division of either the CD56^dim^ or CD56^bright^ NK cells, as judged using by a 5-day CFSE assay (Figure 4G and H). This was in sharp contrast to the potent mitogenic activity of IL-15, with the CD56^bright^ NK cells being more responsive to IL-15 than the CD56^dim^ subset (Figure 4G), consistent with previous data on this cytokine (47). These results suggested that reovirus treatment of PBMC was inducing a number of components associated with proliferation in NK cells, but that the signals delivered were insufficient to initiate proliferation itself.

**Figure 4:**
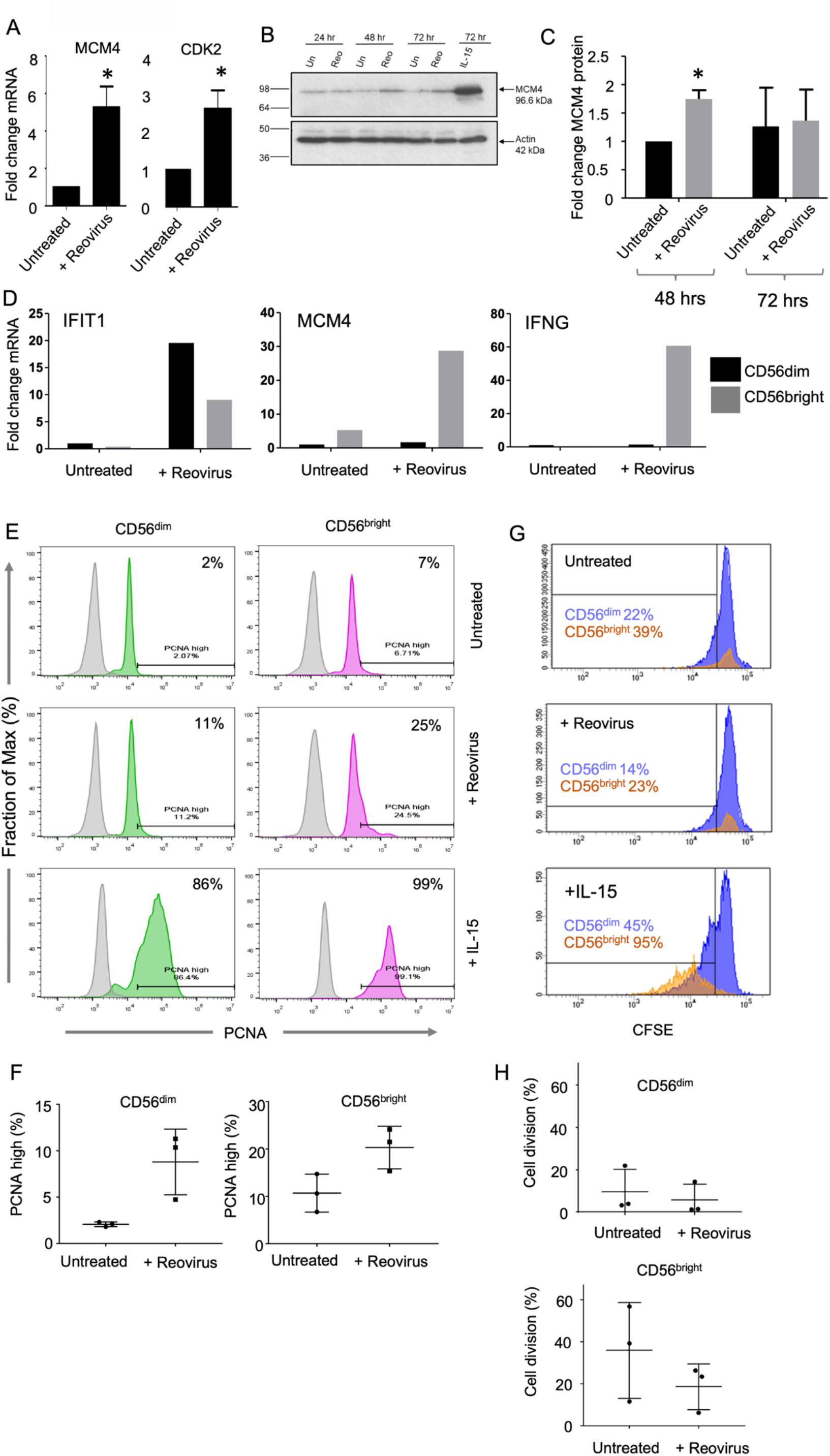
Cell cycle analysis in NK cells following reovirus treatment. **A)** Induction of MCM4 and CDK2 genes in NK cells following reovirus treatment. PBMC were treated with reovirus for 48 hours, NK cells were purified and gene expression determined by qRT-PCR (in 3 or 4 donors respectively). Differences between mean ddCt values were analysed by a paired T test; *p<0.05. **B)** Time course of MCM4 protein expression in NK cells following reovirus treatment. PBMC were cultured with or without 1 MOI reovirus, or with 10 ng/ml IL-15, for either 24, 48 or 72 hours. Protein expression of MCM4 (and β-actin) was analysed by immunoblotting; the data shown is from a single donor. **C)** Quantitation of immunoblotting from three donors, with the fold change in MCM4 expression determined by comparison to β actin and analysed by T test; *p<0.05. **D)** Induction of MCM4, IFNG and IFIT1 genes in CD56^dim^ and CD56^bright^ NK cells following reovirus treatment. Following reovirus treatment of PBMC, CD56^dim^ and CD56^bright^ subsets were purified by cell sorting and gene expression determined by qRT-PCR. Expression is shown in the different treatments relative to the untreated controls, as indicated. These data are from a single donor and are representative of two donors analysed. The cell sorting gates are shown in Supplementary Figure 1C. **E)** PCNA expression in CD56^dim^ and CD56^bright^ NK cells following reovirus treatment. PBMC were cultured with or without reovirus, or with 10 ng/ml IL-15, for 48 hours and PCNA expression analysed by intracellular staining and flow cytometry. Data from a single donor is shown with the PCNA high gate indicated (set as the top 2% of untreated CD56^dim^ cells). The percentage of PCNA high expressing cells is indicated for each treatment. **F)** Mean percentage of PCNA high cells in CD56^dim^ and CD56^bright^ NK cells (determined as in E) from three donors. Differences were tested by paired T test and were not statistically significant. **G)** Cell division following reovirus treatment. PBMC were labelled with CFSE and cultured with or without 1 MOI reovirus, or 10 ng/ml IL-15. After 5 days, CFSE content in CD56^dim^ and CD56^bright^ NK cells was determined by flow cytometry as indicated. The percentage of cells from each subset that fell to the left of the indicated gate and had lost CFSE content due to cell division are indicated. These data are from a single donor. **H)** Proliferation of CD56^dim^ and CD56^bright^ NK cells from the CFSE assay performed in three separate donors (as in G). Differences between treatment means were tested using a paired T test and were not statistically significant.

The induction of the IL15RA gene encoding the high affinity *α*-subunit of the IL-15 receptor (Figure 3E), suggested that reovirus treatment might prime NK cells for subsequent proliferative responses, as shown using mouse models and other viruses (33, 34). We pre-treated PBMC with reovirus for 4, 24 or 48hrs followed by IL-15 for a further 3 days and analysed the cell cycle profile. Addition of IL-15 alone induced S phase, but pre-treatment with reovirus for as little as 4 hours caused a block in cell cycle progression (Figure 5A). Pre-treatment with reovirus blocked the IL-15 mediated proliferation of both the CD56^dim^ and CD56^bright^ subsets, with statistically significant impairment of the more proliferative CD56^bright^ NK cells (Figure 5B). Conditioned media collected from reovirus treated PBMC (filtered to remove virions) and applied to fresh PBMC prior to IL-15 stimulation showed an inhibition of proliferation of the CD56^bright^ NK cell subset that was restored in the presence of blocking antibodies against IFN-I and its receptor, revealing an inhibitory role for reovirus-induced IFN-I in NK cell proliferation (Figure 5C). This was confirmed by stimulating purified NK cells with IL-15 or IL-15 plus IFN-I, showing that IFN-I reduced the mitogenic activity of IL-15, as determined by cell cycle profiling (Figure 5D). We analysed the effects of reovirus pre-treatment on IL-15 mediated induction of DNA proliferation components. Induction of both PCNA and Ki67 were blunted in the presence of reovirus in both the CD56^dim^ and CD56^bright^ NK cells (Figure 5E). Furthermore, immunoblotting showed that pre-treatment with reovirus reduced the induction of MCM4, cyclin B and CDK2 in response to IL-15 (Figure 5F). The ability of reovirus treatment (and IFN-I) to block IL-15 mediated proliferation of NK cells suggested that IL-15 mediated signalling events might be disrupted. We treated PBMC with reovirus for 48 hrs then added IL-15 for 30 minutes and analysed the phosphorylation of STAT5, mTOR and AKT in NK cells using intracellular flow cytometry. The IL-15 mediated phosphorylation of STAT5 and mTOR was unaffected by pre-treatment with reovirus, but IL-15 induced AKT phosphorylation was significantly reduced (Figure 5G). Reovirus pre-treatment also enhanced STAT1 phosphorylation downstream of IL-15, a pathway not typically activated by this cytokine (Figure 5G). Taken together, these data show that reovirus primed NK cells have skewed signalling pathway activation and reduced proliferation in response to IL-15.

**Figure 5:**
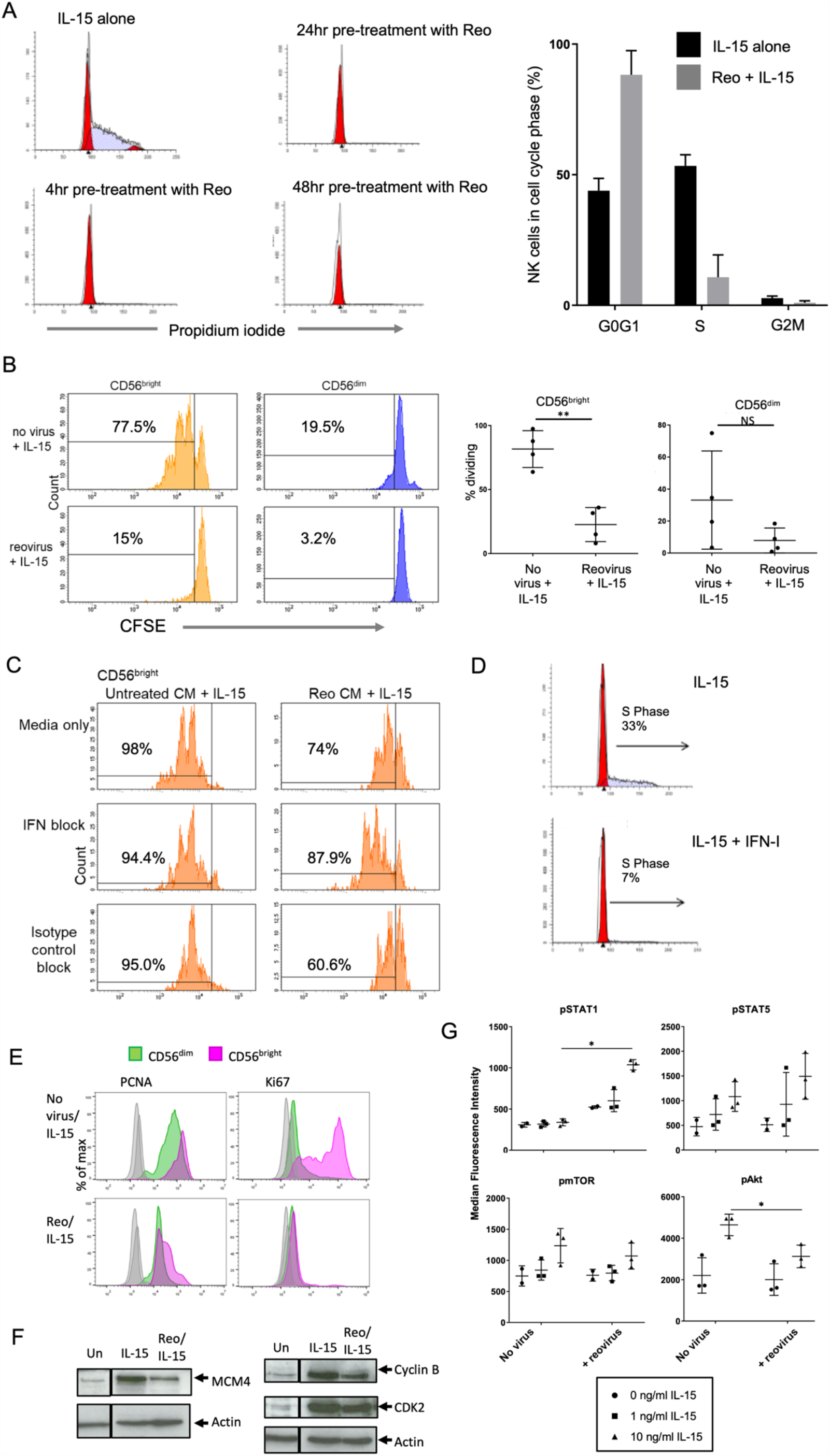
Inhibition of NK cell proliferation following reovirus treatment. **A)** Cell cycle profiling by DNA content. PBMC were treated with either 10ng/ml IL-15 for 3 days, or primed with 1 MOI reovirus for 4, 24 or 48 hours, followed by IL-15 treatment for 3 days. NK cells were purified, fixed and DNA stained with propidium iodide and analysed by flow cytometry. Cell cycle profiles were analysed using Modfit software to quantify the percentage of cells in each compartment, as shown in the graph (collated from three donors). **B)** Reovirus mediated inhibition of IL-15 induced NK cell proliferation. PBMC were labelled with CFSE, primed for 4 hours with 1 MOI reovirus or cultured without virus. 10 ng/ml IL-15 was then added and cells were cultured for a further 5 days. Cell division was assessed by CFSE loss using flow cytometry. Plots from a single donor are shown on the left with the percentage of cells that have proliferated after 5 days indicated. Data from four donors are shown on the right, analysed using a T test; **p<0.01. NS = Not significant. **C)** Inhibition of CD56^bright^ NK cell proliferation by reovirus-induced IFN-I. CFSE labelled PBMC were cultured in conditioned media from unstimulated PBMC (untreated CM) or reovirus treated PBMC (Reo CM), with or without the IFN-blocking antibody cocktail or with a control blocking cocktail. 10 ng/ml IL-15 was added and cells were cultured for 5 days and proliferation analysed by CFSE content of the CD56bright NK cells. The percentage of cells that have proliferated after 5 days are indicated. These data are from a single donor, representative of 3 donors. **D)** IFN-I blocks IL-15 induced S phase. Purified NK cells were cultured with IL-15 or IL-15+IFN-I (all at 100 ng/ml) for 3 days and S phase assessed by propidium iodide staining, as shown in panel A). One donor, representative of three donors tested. **E)** Reovirus treatment blocks IL-15 induced expression of proliferation markers. PBMC were cultured for 4 hours alone (no virus) or primed with 1 MOI reovirus (Reo). 10 ng/ml IL-15 was then added directly to all samples for 48 hours. Expression of PCNA and Ki67 was determined using intracellular staining and flow cytometry in the CD56^dim^ (green) and CD56^bright^ (purple) NK cells. These data are representative of 3 donors tested. **F)** Reovirus treatment blocks IL-15 induced expression of cell cycle mediators. PBMC were cultured as in E), for 3 days. Total NK cells were isolated and MCM4, cyclin B and CDK2 analysed by immunoblotting along with β-actin as a loading control. The blot image has been cut between the unstimulated control and cytokine treated lanes as shown by the boxing. These data are representative of 3 donors tested. **G)** Modulation of IL-15 mediated signalling by reovirus treatment. PBMC were cultured for 48 hours alone (no virus) or primed with 1 MOI reovirus (+ reovirus). After 48 hours, 0, 1 or 10 ng/ml IL-15 was added (as indicated) for 30 minutes. Phospho-STAT1, STAT3, STAT5, mTOR and Akt were analysed by intracellular staining and flow cytometry, gating on the NK cell population. Graphs show median fluorescence intensities (MFI) for 2 or 3 separate donors. Data were analysed by two-way repeated measures ANOVA. When the effect of virus was statistically significant, a post-hoc Sidak multiple comparison test was applied to identify statistically significant differences between “no virus” and “reovirus” MFI values; *p<0.05.

Amongst the genes downregulated in NK cells following reovirus treatment of PBMC was the sphingosine-1-phosphate receptor, S1PR1 (∼5-fold reduction; Figure 3A), a regulator of lymphocyte trafficking (48). For mouse B and T lymphocytes, IFN-I induced CD69 expression decreases sphingosine-1-phosphate receptor activity, contributing to the retention of these cells in lymph nodes (49). We analysed changes in S1PR1 gene expression alongside CCR7, a chemokine receptor implicated in lymph node homing (50). Reovirus treatment was associated with reduced S1PR1 and increased CCR7 mRNA (Figure 6A). Coupled with the IFN-I induced expression of CD69 both *in vitro* (Figure 3A and Supplementary Figure S3) and *in vivo* (21), this phenotype suggested that reovirus treatment might alter the tissue distribution of NK cells during therapy.

**Figure 6:**
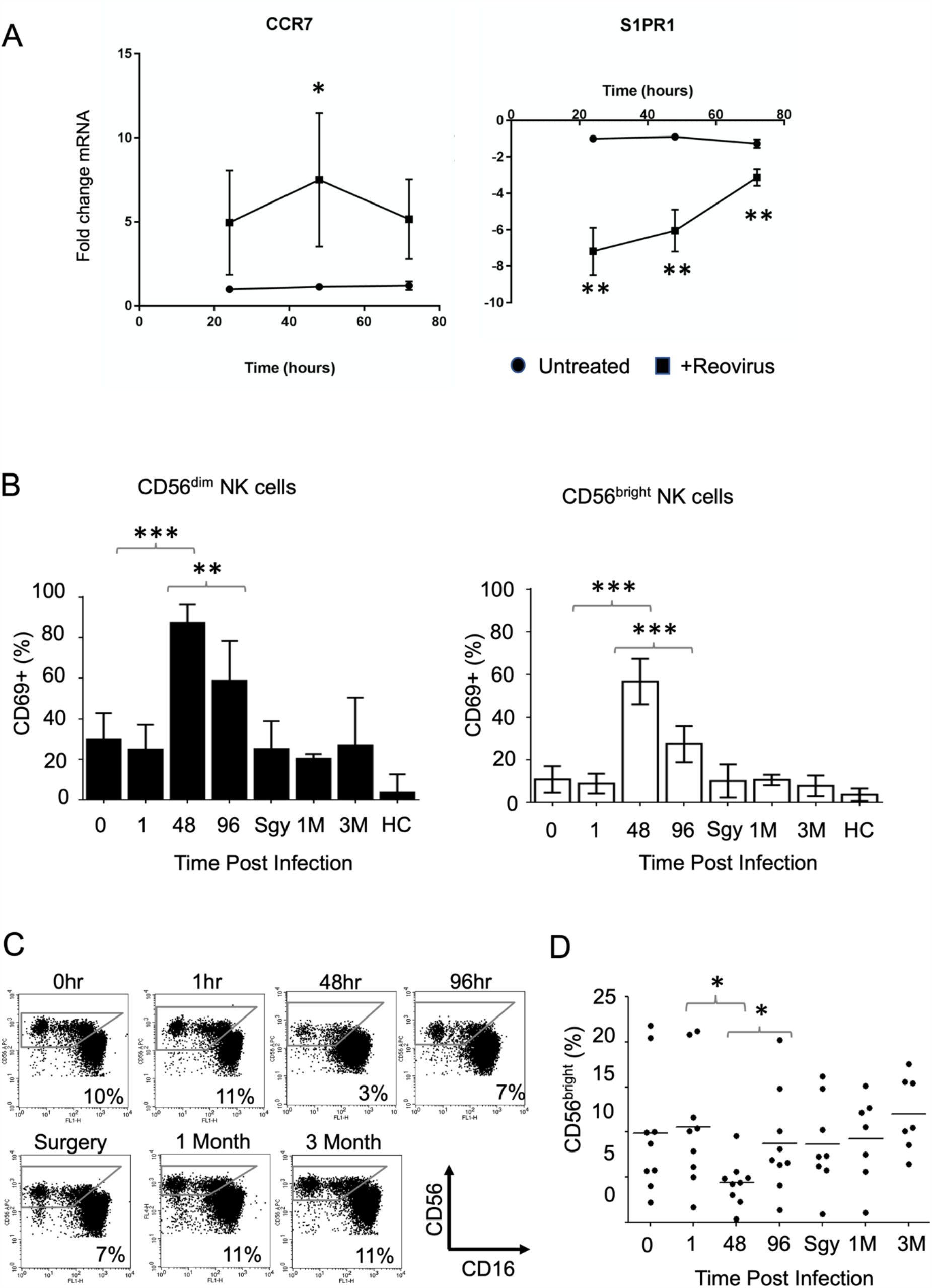
Reovirus treatment alters human NK cell subset distribution *in vivo*. **A)** Expression of CCR7 and S1PR1 genes in NK cells following reovirus treatment *in vitro*. PBMC were treated with reovirus for 24, 48 and 72 hours, NK cells were isolated and gene expression assessed using qRT-PCR. The data show the fold change in expression in reovirus treated cells compared to untreated, indicating the mean and standard deviation from three donors. The effect of time and treatment on mean ddCt values was tested by two-way repeated measures ANOVA, followed by Sidak multiple comparisons test; *p<0.05, **p<0.01 **B)** NK cell CD69 expression following reovirus treatment of patients. Nine cancer patients were treated with intravenous reovirus at time zero with blood samples taken immediately prior to infusion (0hr) and at 1hr, 48hr and 96 post-infusion. Patients had another blood sample taken immediately prior to surgery (Sgy) 1-4 weeks after infusion and additional samples 1 month (1M) and 3 months (3M) later. Samples were also taken from untreated healthy controls (HC); full details of the patients, controls and treatment scheduling have been published previously (20). Blood samples were used to assess NK cell activation by cell surface CD69 expression on the CD56dim (left panel) and CD56bright (right panel) NK cell subsets. Statistically significant changes in CD69 expression, determined using a Wilcoxon Rank test, are indicated; **p<0.01, ***p<0.001. Note that ten patients in total were treated in the trial (20), but one patient had samples taken at different times, preventing their inclusion in this analysis. **C)** Loss of CD56^bright^ NK cells from the circulation at the peak of NK cell activation. The percentage of CD56^bright^ NK cells (defined as CD56^bright^, CD16^neg/low^, CD3^neg^) was determined as a fraction of the total NK cell population during reovirus treatment; this panel shows one presentative patient. **D)** Percentage of CD56^bright^ NK cells as a fraction of the total NK cell population in nine reovirus treated patients. Statistically significant changes in the size of the CD56^bright^ NK cell population were determined using a Wilcoxon Rank test; *p<0.05.

Using blood samples from reovirus-treated patients, we previously showed that the peak of NK cell CD69 expression coincided with the peak IFN-I response (21). We analysed CD69 expression in these patients in more detail, gating separately on CD56^bright^ and CD56^dim^ NK cell populations; CD69 expression was induced on both subsets with similar kinetics. Importantly, CD69 expression on both CD56^bright^ and CD56^dim^ NK cells was significantly enhanced from 1hr to 48hrs post-treatment and significantly decreased from 48hrs to 96 hrs post-treatment (Figure 6B) with peak CD69 expression in both subsets matching the peak IFN-I response shown previously (21). We analysed the relative abundance of the two NK cell subsets over the treatment course and found that, as a proportion of the total NK cells, the CD56^bright^ NK cells were significantly reduced in the blood 48 hrs post-treatment (Figure 6C and D). Thus, for the CD56^bright^ NK cell subset, the expression of cell surface CD69 is inversely related to the abundance of these cells in the blood and as CD69 expression dropped 96 hrs post-treatment, the CD56^bright^ NK cells showed significantly increased abundance in the blood (Figure 6C and D). These results reveal that oncolytic reovirus treatment transiently alters the tissue distribution of human NK cell subsets *in vivo*.

## 4. Discussion

We have analysed the mechanisms by which oncolytic reovirus modulates human NK cell activity. A previous clinical trial of intravenous reovirus in cancer patients identified a peak of NK cell activation approximately 48hrs post-infusion. This was co-incident with the maximal IFN-I response as determined by ISG expression in leucocytes. Not surprisingly, NK cell activation in these patients bore the hallmarks of an IFN-I response (21). Related *ex vivo* studies have shown that reovirus triggers IFN-I production by monocytes and that reovirus-mediated enhancement of NK cell cytotoxicity is both monocyte and IFN-I dependent (22). The kinetics of NK cell activation in these oncolytic reovirus-treated cancer patients resemble those found in mouse models, as well as human studies (2, 51–55). The results presented here demonstrate that reovirus-induced IFN-I acts directly on NK cells, inducing STAT1 phosphorylation and a characteristic transcriptional profile. The ISGs expressed by NK cells following reovirus treatment include many that interfere with various stages of viral entry, replication and egress (56, 57). In addition, analysis of the interferome database coupled with prior studies confirm that components of the NK cell cytotoxic machinery are induced by IFN-I (33, 58–60). These results show that NK cells respond to IFN-I in two ways; like other cells they increase their own anti-viral defence mechanisms and, in addition, they increase expression of cytotoxic components to eliminate infected cells. This allows NK cells to destroy infected cells whilst minimising their own infection.

In mouse models, IFN-I induces IL-15 production by DC which then activates NK cells (33, 34) and for human NK cells, enhancement of reovirus-mediated NK cell cytotoxicity is IL-15 dependent (61). Although not as striking as the IFN-I/STAT1 response, we detected a low (but statistically significant) level of STAT5 phosphorylation in the CD56^bright^ NK cells following reovirus treatment. Furthermore, reovirus treatment induced several genes encoding cell cycle functions in NK cells. Analysis of the regulation of these genes using the interferome database suggests that they are not direct targets of IFN-I, but are more likely induced by other cytokines, with IL-15 being a prime candidate (47, 62). This is consistent with data from mouse models showing that IL-15 blockade following IFN-I treatment inhibits NK cell proliferation, but not cytotoxicity (33).

Although reovirus treatment induced numerous cell cycle genes and their protein products, we did not detect the induction of NK cell proliferation, even when prolonging our assays for up to five days post-reovirus treatment. We considered the possibility that reovirus and IFN-I were priming NK cells for subsequent proliferation (e.g. by inducing the expression of IL15 and IL15RA genes), or that the weak STAT5 phosphorylation we detected in CD56^bright^ NK cells was simply reflective of low levels of IL-15 in these assays. However, blocking and reconstitution experiments showed that reovirus treatment blocks IL-15 mediated NK cell proliferation in an IFN-I dependent manner. This block on proliferation is not surprising; the pro-cytotoxic, but anti-proliferative action of interferon on lymphocytes is long established (63, 64) and, more recently, mouse NK cells lacking a functional IFN-I receptor (*Ifnar*^-/-^) were shown to exhibit enhanced proliferation following MCMV infection (65). However, IFN-I mediated inhibition of proliferation is paradoxical given that NK cell expansion occurs after viral infection in both mouse models and humans (65–70). However, meaningful comparisons of the kinetics of IFN-I responses and NK cell proliferation in these diverse systems are difficult to make. It seems likely that NK cell proliferation occurs once the peak of the IFN-I response has subsided; in the reovirus clinical trial, NK cell expansions were only detectable after the peak of NK cell activation and ISG expression (21). Interestingly, for T cells, IFN-I can have both pro-and anti-proliferative functions dependent upon whether it signals before or after TCR signalling. This dichotomous activity is regulated by the balance of the pro-proliferative STAT4 versus anti-proliferative STAT1 in the pre- or post-TCR stimulated cell (71, 72). Furthermore, in some tumour cells, IFN-I signalling results in the induction of the cyclin-dependent kinase (CDK) inhibitor, p21^WAF1^ (73, 74); such activity preceding TCR signalling (or in NK cells, IL-15 receptor engagement) might prevent proliferation. We did not detect significant induction of CDKN1A (p21^WAF1^), CDKN1B (p27^KIP1^) or STAT4 mRNA in NK cells from reovirus treated PBMC, but STAT1 mRNA was highly induced. Furthermore, PI3K-AKT signalling is essential for mouse NK cell proliferation (75) and IL-15 induced AKT phosphorylation was reduced in NK cells pre-treated with reovirus; reduced AKT phosphorylation may also antagonise nuclear exclusion of p27^KIP1^, favouring inhibition of proliferation (76, 77).

Gene expression changes suggestive of an altered NK cell trafficking phenotype prompted us to analyse NK cell subset distribution in reovirus treated patients. These patients showed the selective loss of the CD56^bright^ NK cell subset from the blood 48 hrs post-infusion. This timepoint coincides with the peak IFN-I response following reovirus delivery *in vivo* (21). The egress of NK cells from lymph nodes in response to S1P differs from B and T lymphocytes, with S1PR5 as well as S1PR1 regulating activity (48, 49, 78, 79); only S1PR1 mRNA met our statistical criteria for differential expression following reovirus treatment. The CD56^bright^ NK cell subset is highly enriched in SLT and it appears likely that reovirus treatment reduces S1P-mediated egress from SLT in an IFN-I dependent manner. Reovirus treatment induced IFNG expression in CD56^bright^ NK cells and, by analogy with mouse models, the transient retention of IFN-γ expressing CD56^bright^ NK cells in the SLT will favour Th1 differentiation and more effective cytotoxic T cell responses (80). However, S1P gradients also regulate NK cell trafficking to inflamed tissues (79) and, since we were only able to sample blood from these patients, we cannot rule out the possibility that the CD56^bright^ NK cells lost from the circulation have relocated to the tumour in the liver, where we previously showed reovirus to be replicating (28).

The anti-tumour action of OV stems from the combination of direct tumour lysis and the induction of anti-tumour immunity, and is dependent upon NK cell activity (13, 17–20). Our results show that OV treatment regulates both NK cell cytotoxicity and the immunomodulatory activity of CD56^bright^ NK cells, the latter contributing to the induction of adaptive anti-tumour responses. Our results also show that OV-mediated NK cell responses are in part driven by the direct action of IFN-I on NK cells which activates effector functions, but inhibits IL-15 mediated proliferative responses. These OV-mediated effects presumably reflect the spectrum of human NK cell responses seen during normal viral infection.

## 5. Acknowledgements

This work was supported by a Bramall PhD Scholarship (to MW) and grant awards from the Medical Research Foundation and Cancer Research UK.

## 6. Author contributions and declarations

Conceived and designed study: MW, AAM, LFW, GPC.

Performed experimental work: MW, LFW, SLP, EBW

Performed bioinformatics analysis: MW, APD, GPC

Designed, implemented and analysed clinical trial: AAM, MC, YES, TDH, GPC

Wrote paper: MW and GPC with input from all authors

Pelareorep for the clinical trial and for laboratory studies was provided by Oncolytics Biotech. Matt Coffey is an employee of Oncolytics Biotech with shares and stock options. Alan Melcher has previously received research grant funding from Oncolytics Biotech.

## 9. Legends to Supplementary Figures and Tables

**Supplementary Tables 1 and 2:**

List of antibodies and PCR primers used in this study.

**Supplementary Table 3**.

Gene expression profiling of NK cells following reovirus treatment. The table shows the 1742 differentially expressed genes (FDR<0.05, fold change >1.5X) that were either upregulated (939 genes; highlighted in green) or downregulated (803 genes; highlighted in red) in NK cells from reovirus treated PBMC compared to untreated PBMC. Data from this table were used to construct Figures 3A-E of the main manuscript.

**Supplementary Table 4**.

Pathways represented by differentially expressed genes identified using gene set enrichment analysis. The 1742 genes (listed in Supplementary Table 3) were inputted into the Enrichr tool (28, 29; https://amp.pharm.mssm.edu/Enrichr/) and enriched pathways identified using the Reactome 2016 database. The table shows the output from Enrichr, listing the significantly enriched pathways (adjusted p<0.05), and those genes from the DEGs that are assigned to these pathways. Data from this table was used to construct Figure 3B of the main manuscript.

**Supplementary Table 5**.

Transcription factors predicted to be associated with the differentially expressed genes identified using gene set enrichment analysis. The 1742 genes (listed in Supplementary Table 3) were inputted into the Enrichr tool (28, 29; https://amp.pharm.mssm.edu/Enrichr/) and candidate transcription factors analysed from the Encode and ChEA Consensus TFs from ChIP-X database. The table shows the output from Enrichr, listing the significantly enriched transcription factors (adjusted p<0.05) and those genes from the DEGs that are associated with these transcription factors. Data from this table was used to construct Figure 3C of the main manuscript.

**Supplementary Figure S1**.

**Gating strategies to separate CD56**^**dim**^ **and CD56**^**bright**^ **NK cells**

**A)** Representative gating of CD56^dim^ and CD56^bright^ NK cells for Phos-Flow, surface and intracellular staining experiments.

**B)** Gating of CD56^dim^CD16+ and CD56^bright^ CD16-NK cells after 5 day CFSE culture experiments.

**C)** Gating used in cell sorting of CD56^dim^CD16+ and CD56^bright^ CD16-NK, after magnetic isolation of NK cells from PBMC.

**Supplementary Figure S2**

**Intracellular staining and flow cytometry to detect STAT phosphorylation**

PBMC were treated with 100 IU/ml IFNα, 10 ng/ml IL-12 or 50 IU/ml IL-15 for 1 hour. NK cells within the PBMC (detected using cell surface staining as shown in Supplementary Figure 1A) were evaluated for levels of phosphorylated STAT1, STAT4 and STAT5 by intracellular flow cytometry. Histograms show isotype control or phosphorylated protein staining in total NK cells, representative of 3 donors.

**Supplementary Figure S3**

**CD69 expression following reovirus treatment**

**A)** PBMC from five donors were treated with reovirus for 48hrs *in vitro* and the NK cells purified by magnetic immunoselection. Activation of the NK cells was verified by analysis of CD69 expression in all five donors. Four donors (1, 2, 4 and 5) were analysed using a FITC-conjugated antibody (left hand panels) and one (donor 3) was analysed using a PE-conjugated antibody (right hand panels). These NK cells were used as a source of mRNA for gene expression profiling.

**B)** Expression of CD69 on CD56^dim^ and CD56^bright^ NK cells. The values indicate the percentage of cells expressing CD69 (or not) for both CD56^bright^ NK cells (top two quadrants) and CD56^dim^ NK cells (bottom two quadrants) in both untreated PBMC and PBMC treated with reovirus for 48 hrs

**Supplementary Figure S4**

**Reovirus and IFN-I regulate NK cell cytotoxicity**.

**A)** PBMC were cultured without stimulation (untreated) or with reovirus (MOI 1) for 48 hours, followed by co-culture with K562 target cells. Degranulation of NK cells determined by CD107a expression on CD56+ NK cells. Plots are representative of data obtained from 2 donors.

**B)** IFN-I mediated induction of granzyme B. Purified NK cells were left untreated or stimulated with 10ng/ml IL-15 or increasing amounts of IFN-I (50 IU, 100 IU and 200 IU) for 48 hrs. Granzyme B expression was determined by immunoblotting, using actin as a control.

**C)** Purified NK cells were left untreated (Un) or stimulated with 10ng/ml IL-15 or 100 IU of IFN-I for 48 hrs and used in a degranulation assay (as in panel A) against K562 cells. The experiment was performed in triplicate and analysed using a Students T test.

**Supplementary Figure S5**

**Expression of CCNB1 mRNA in response to reovirus**.

PBMC were untreated (un) or treated with reovirus for 48 hours. NK cells were then isolated and CCNB1 gene expression analysed by qRT-PCR. Mean relative expression and standard deviation from three donors. Differences between mean ddCt values were tested by paired T test and were not significant.

**Supplementary Figure S6**

**Ki67 protein expression is not upregulated with reovirus treatment**.

PBMC were cultured with (Reo) or without (Un) 1 MOI reovirus, or with 10 ng/ml IL-15, for 48 hours. Protein expression of Ki67 was analysed in CD56^dim^ and CD56^bright^ NK cell populations by intracellular flow cytometry.

**A)** Representative histograms of ki67 expression from 1 of 3 donors. Untreated cells were used to set a gate for ki67 low cells (98%) or ki67 high expressing cells (2%). This gate was used to determine ki67 expression in the Reo and IL-15 treated samples in both CD56^dim^ and CD56^bright^ NK cell subsets.

**B)** Data from three donors, showing the percentage of ki67 high expressing NK cells in untreated and reovirus treated PBMC. Differences between mean percentage values were analysed using a paired T test.

