## Supplementary Figures S1-S6 for "Oncolytic virus treatment differentially affects the CD56^dim^ and CD56^bright^ NK cell subsets *in vivo* and regulates a spectrum of human NK cell activity"

**Supplementary Figure S1**

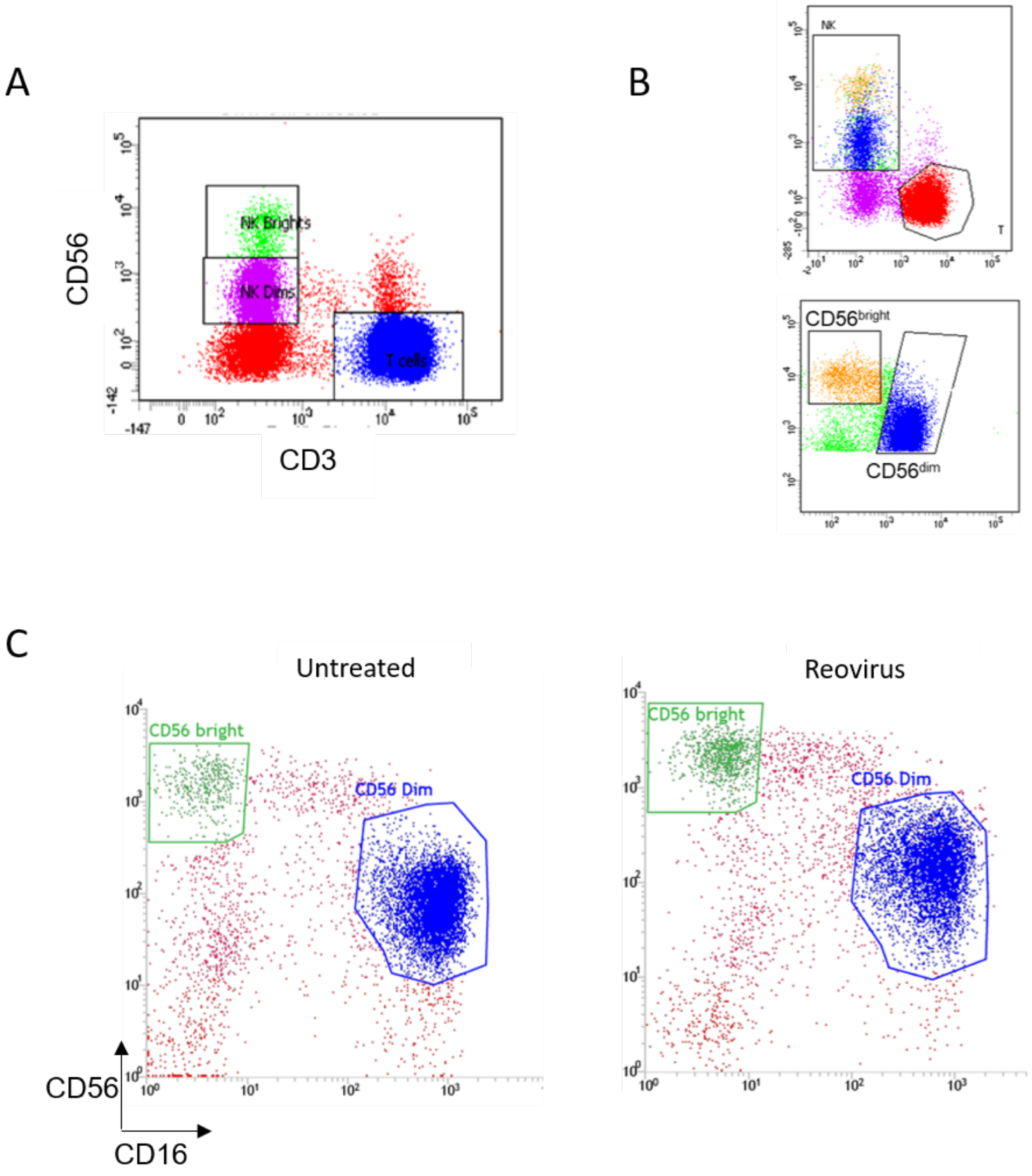

**Supplementary Figure S2**

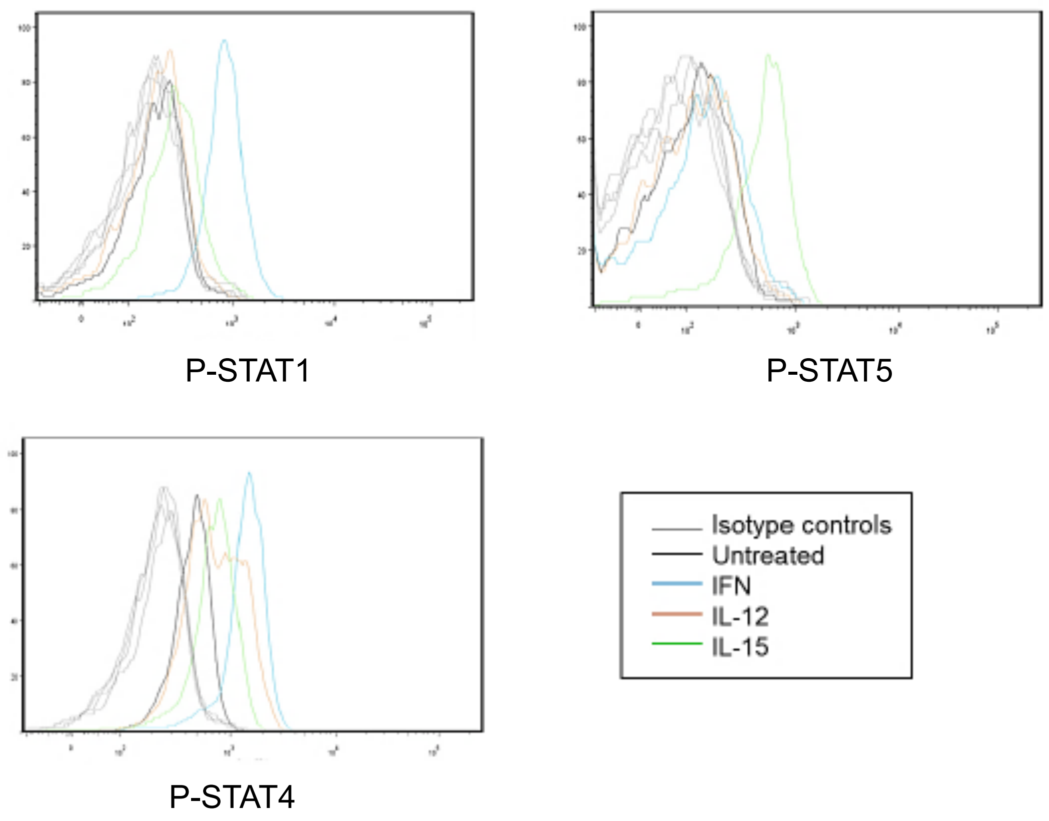

**Supplementary Figure S3**

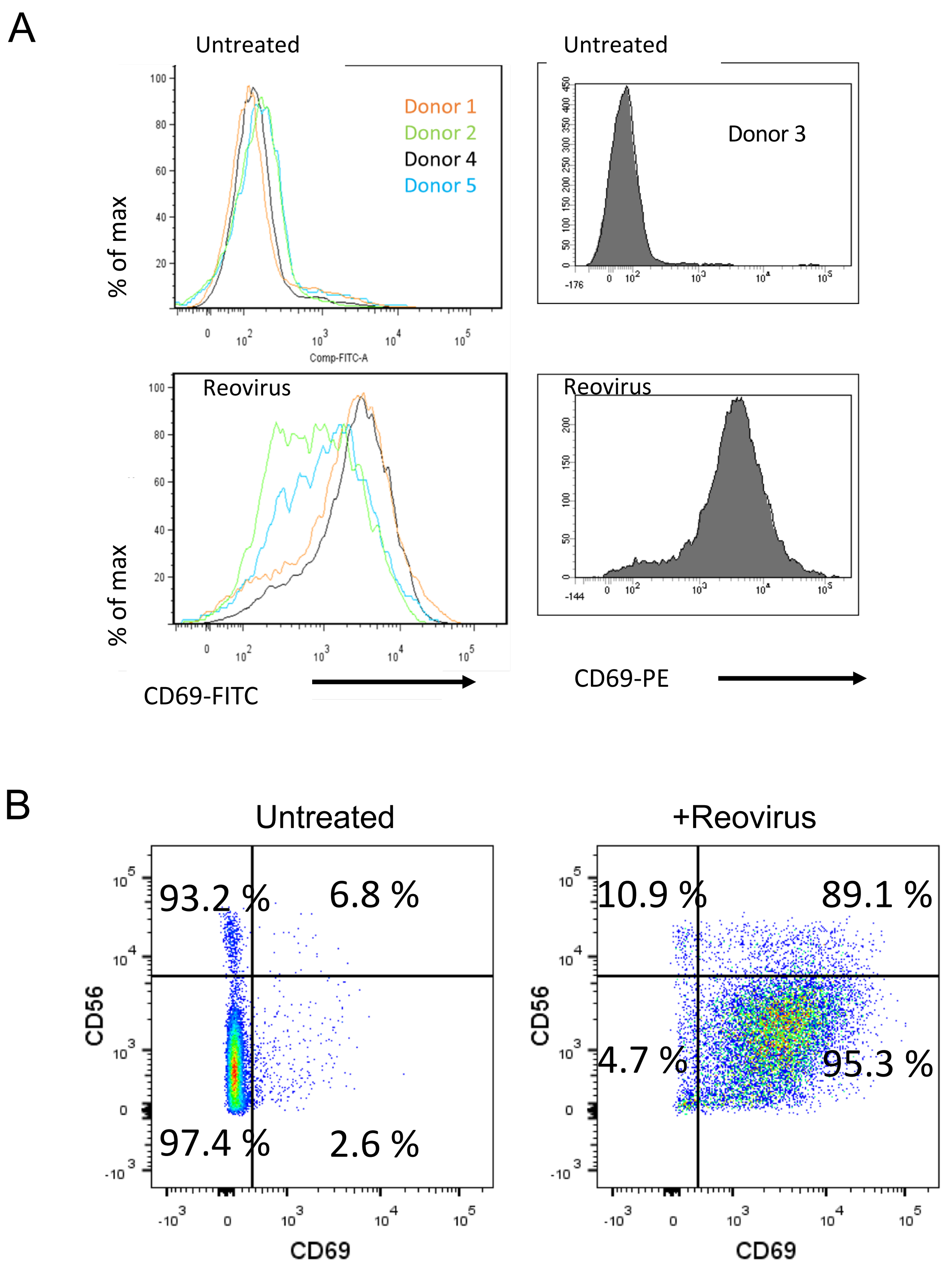

### Supplementary Figure S4

A

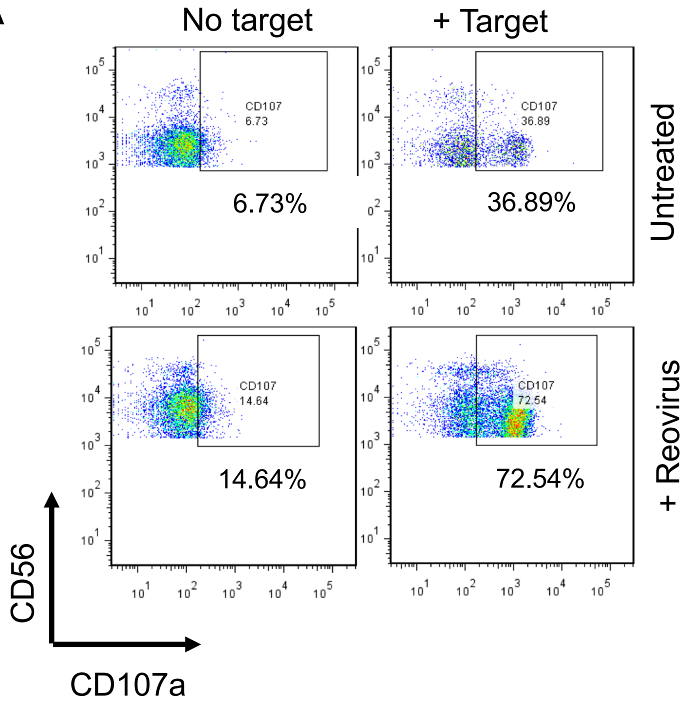


B

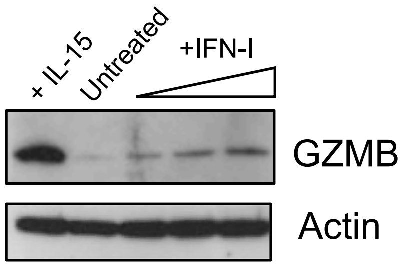


C

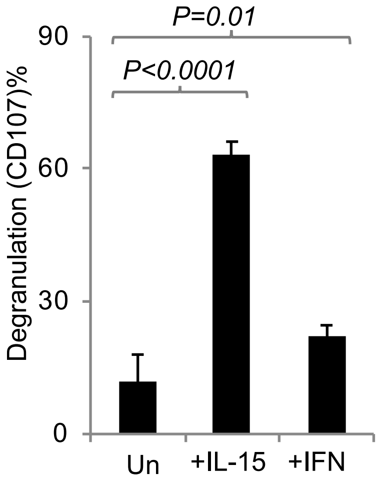

**Supplementary Figure S5**

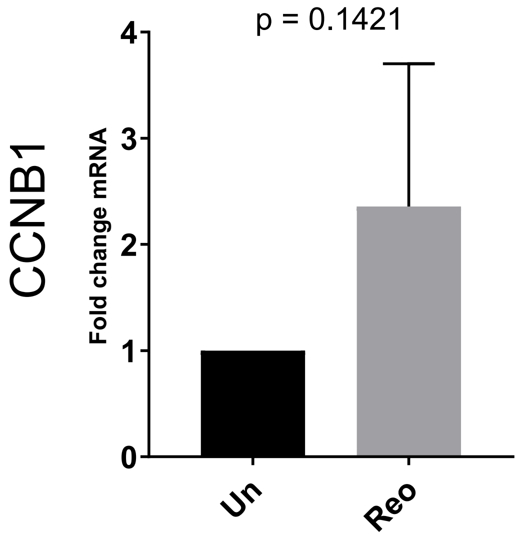

**Supplementary Figure S6**

A

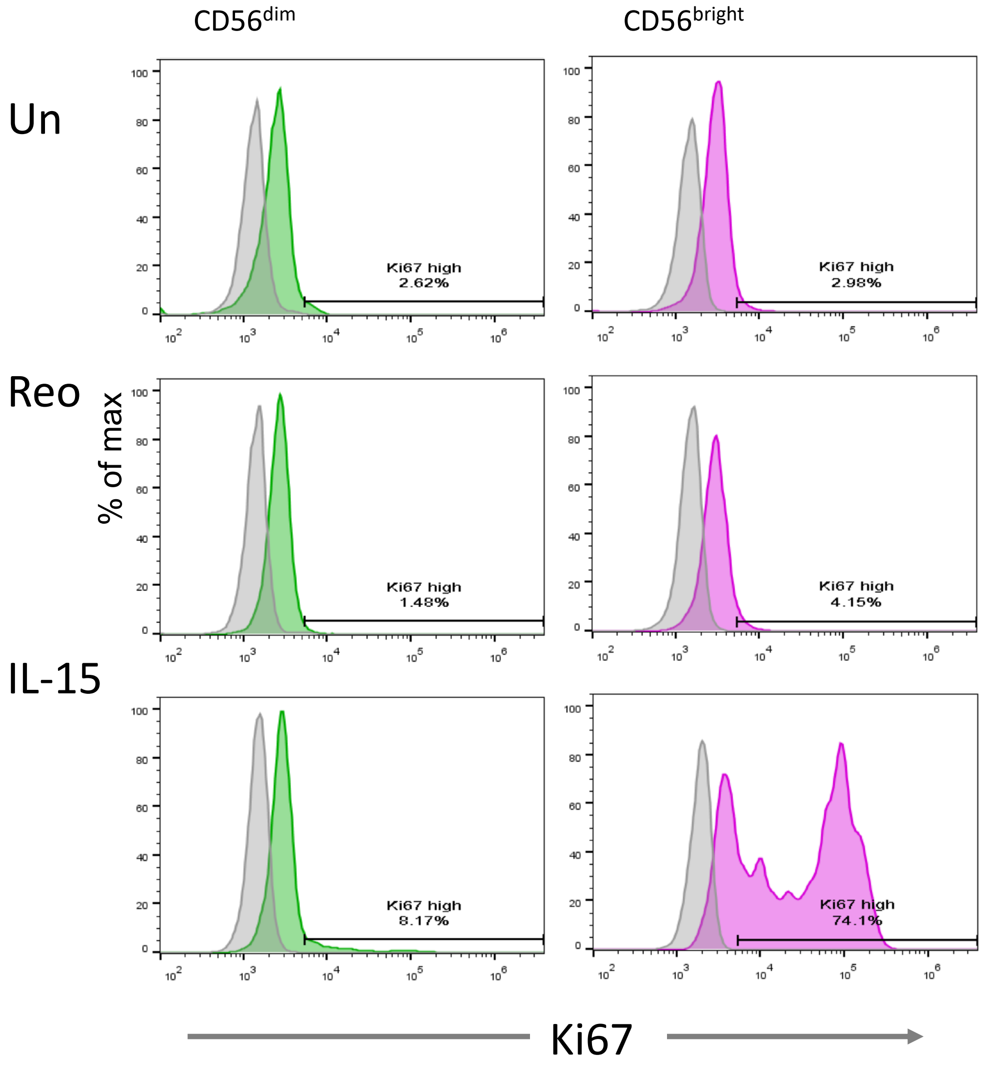


B

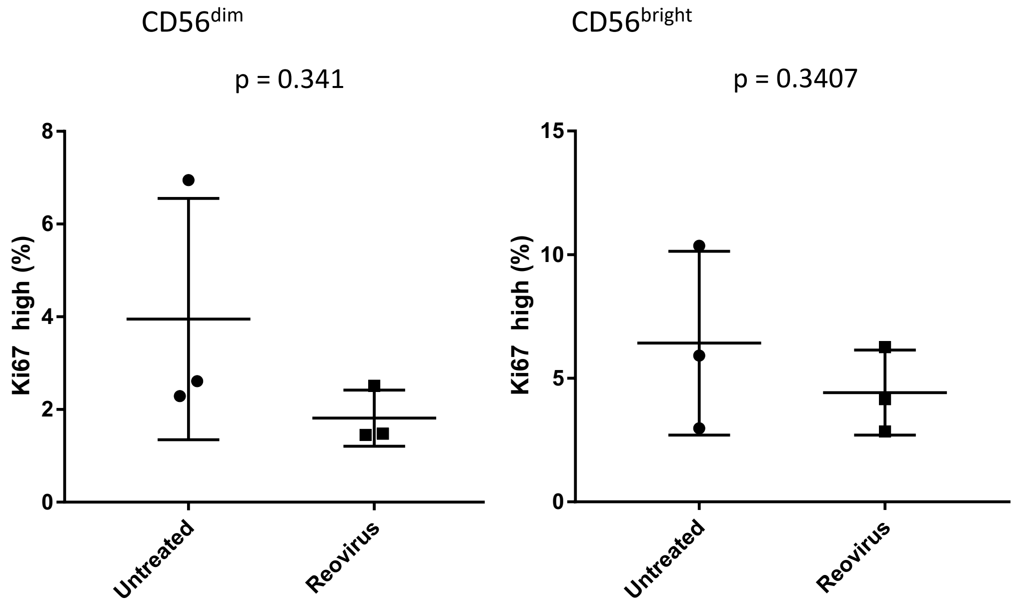
