## Supplementary Tables 1 and 2 for "Oncolytic virus treatment differentially affects the CD56^dim^ and CD56^bright^ NK cell subsets *in vivo* and regulates a spectrum of human NK cell activity"

### Table 1: Antibodies used in this study:

| **Immunoblotting** | | | | |
|---|---|---|---|---|
| **Antigen** | **Species** | **Clone/ID** | | **Manufacturer** |
| STAT1 | mouse | 1 | | BD Biosciences |
| pSTAT1 (pY701) | mouse | 4a | | BD Biosciences |
| STAT4 | mouse | Clone 8 | | BD Biosciences |
| pSTAT4 (pY693) | mouse | 38 | | BD Biosciences |
| STAT5 | rabbit | 3H7 | | Cell Signalling Technology |
| pSTAT5 (Y694) | rabbit | C11C5 | | Cell Signalling Technology |
| MCM4 | rabbit | Ab4459 | | Abcam |
| Granzyme B | mouse | 2C5/F5 | | BD Biosciences |
| β-Actin | mouse | AC-15 | | Sigma |
| **Flow cytometry** | | | | |
| **Antigen** | **Conjugate** | **Clone/ID** | **Isotype** | **Manufacturer** |
| CD56 | PE-Vio770 | REA196 | Human IgG1 | Miltenyi |
| CD56 | APC | AF12-7H3 | IgG1 | Miltenyi |
| CD3 | FITC | UCHT1 | IgG1 | BD Biosciences |
| CD3 | BV421 | UCHT1 | IgG1 | BD Biosciences |
| CD69 | BV421 | FN50 | IgG1 | BioLegend |
| CD69 | FITC | FN50 | IgG1 | BioLegend |
| CD69 | PE | FN50 | IgG1 | BioLegend |
| CD317 (Tetherin) | PE | REA202 | Human IgG1 | Miltenyi |
| pY701 STAT1 | APC | REA345 | Human IgG1 | Miltenyi |
| pY693 STAT4 | PE | 38/p-Stat4 | IgG2b | BD Biosciences |
| pY694 STAT5 | PerCP-Cy5.5 | 47/Stat5(pY694) | IgG1 | BD Biosciences |
| Granzyme B | PE | GB11 | IgG1 | Thermo Fisher Scientific |
| CD253 (TRAIL) | APC | RIK-2.1 | IgG1 | Miltenyi |
| CD16 | BV421 | 3G8 | IgG1 | BioLegend |
| CD16 | BUV395 | 3G8 | IgG1 | BD Biosciences |
| PCNA | PE | PC10 | IgG2a | BD Biosciences |
| Ki67 | PE | REA183 | Human IgG1 | Miltenyi |
| pY694 STAT5 | BV421 | 47/Stat5(pY694) | IgG1 | BD Biosciences |
| pS2448 mTOR | PE | O21-404 | IgG1 | BD Biosciences |
| pS473 AKT | Alexa Fluor 488 | M89-61 | IgG1 | BD Biosciences |

| | | | | |
|---|---|---|---|---|
| CD107a | PE | H4A3 | IgG1 | BD Biosciences |
| **Isotype Controls** | | | | |
| | BV421 | MOPC-21 | IgG1 | BioLegend |
| | FITC | MOPC-21 | IgG1 | BD Biosciences |
| | PE | REA293 | Human IgG1 | Miltenyi |
| | APC | REA293 | Human IgG1 | Miltenyi |
| | PE | 27-35 | IgG2b | BD Biosciences |
| | PerCP-Cy5.5 | MOPC-21 | IgG1 | BD Biosciences |
| | PE | MOPC-31C | IgG1 | BD Biosciences |
| | VioBlue | REA293 | Human IgG1 | Miltenyi |
| | PE | MOPC-173 | IgG2a | BD Biosciences |
| | BV421 | X40 | IgG1 | BD Biosciences |
| | Alexa Fluor 488 | MOPC-21 | IgG1 | BD Biosciences |

#### Table 2: qRT-PCR primers

##### 2.1 SYBR Green method

CDK2 F: ATGGATGCCTCTGCTCTCACTG R: CCCGATGAGAATGGCAGAAAGC

CCNA2 F: CTCTACACAGTCACGGGACAAAG R: CTGTGGTGCTTTGAGGTAGGTC

CCNB1 F: GACCTGTGTCAGGCTTTCTCTG R: GGTATTTTGGTCTGACTGCTTGC

##### 2.2. Taqman method

ABL1: Applied Biosystems Hs01104728_m1

CCR7: Applied Biosystems Hs01013469_m1

S1PR1: Applied Biosystems Hs01922614_s1

MCM4: Applied Biosystems Hs00907398_m1

IFNG: Applied Biosystems Hs00989291_m1
